# Spleen tyrosine kinase inhibition mitigates hemin-induced thromboinflammation in the lung and kidney of sickle cell mice

**DOI:** 10.1101/2024.05.04.592537

**Authors:** Juma El-Awaisi, Gina Perrella, Nicolas Mayor, Veronika Tinkova, Simon J Cleary, Beata Grygielska, Steve P Watson, Jordan D Dimitrov, Alexander Brill, Phillip LR Nicolson, Dean Kavanagh, Neena Kalia, Julie Rayes

## Abstract

Sickle cell disease (SCD) leads to hemolytic anemia, vaso-occlusive crisis (VOC), hypoperfusion, and progressive organ damage. Hemin, released during hemolysis in SCD, induces platelet activation through CLEC-2, endothelial activation through TLR4, neutrophil adhesion and NETosis, all of which are regulated by spleen tyrosine kinase (Syk). In this study, we assessed neutrophil and platelet recruitment to the pulmonary, renal, splenic, and hepatic microvasculature in control and SCD mice following hemin injection and the effect of Syk inhibition on cell recruitment and organ perfusion. Compared to controls, SCD mice exhibited higher baseline neutrophil and platelet recruitment to the lungs without alterations in lung perfusion as measured by laser speckle contrast imaging. Injection of hemin increased cell recruitment to the pulmonary and renal vasculature with a concomitant reduction in organ perfusion. However, hemin injection did not change cell recruitment or organ perfusion in the spleen and liver, both of which were altered at baseline in SCD mice. Pretreatment of SCD mice with the Syk inhibitor BI-1002494 mitigated baseline and hemin-induced neutrophil and platelet adhesion in the pulmonary and renal microvasculature, with a corresponding normalization of perfusion. Syk regulates vascular integrity in the lung of SCD mice; whilst high concentrations of BI-1002494 increased bleeding, lowering drug concentrations preserved the inhibitory effect on platelet and neutrophil recruitment and lung perfusion and protected from bleeding complications. These data substantiate Syk as a mediator of vascular thrombo-inflammation and hypoperfusion in the lung and kidney of SCD and provide a rationale for pharmacological inhibition as a therapeutic strategy.

## Introduction

Sickle cell dproisease (SCD) is an inherited severe hemoglobinopathy caused by a mutation on the beta hemoglobin gene, leading to the formation of abnormal hemoglobin S (HbS). Upon deoxygenation, HbS undergoes polymerization, altering the morphology and biomechanical properties of red blood cells (RBCs), inducing sickling and increasing their adhesion ^1^. Clinical manifestations of SCD include chronic hemolytic anemia, episodic and painful vaso-occlusive crises (VOC) that compromise tissue perfusion and lead to progressive end-organ damage ^2,3,4^. Affected individuals suffer an elevated risk of developing acute chest syndrome (ACS), a potentially fatal vaso-occlusive complication within the pulmonary circulation, which is one of the leading causes of hospitalization and premature mortality. High levels of inflammatory markers such as interleukin-6, and platelet-derived thrombospondin-1 and CD40L are associated with ACS ^5,6^. Although specific triggers that result in acute VOC episodes remain poorly defined, contributory factors such as physiological stress ^7^, cold temperature ^8^, hemolysis ^9^, hypoxemia and heightened inflammatory states have been shown to trigger VOC. Hydroxyurea, which increases fetal hemoglobin, remains the standard of care in patients with SCD. It reduces the frequency of painful crises in children and adults and decreases infections in children ^10,11^. However, neutrophils, the main drivers of lung injury ^12,13^, remain active in hydroxyurea-treated patients ^14^. Voxelotor, an inhibitor for hemoglobin S polymerization, alone or in combination with hydroxyurea, was introduced to treat patients over 12 years old but lacks data supporting meaningful improvement in important clinical endpoints. The phase 3 randomized trial of voxelotor in SCD showed increased hemoglobin levels and reduced markers of hemolysis, but only a trend of reduced VOC incidence was observed ^15^. More recently, two CRISPR-Cas9 treatments aiming to increase fetal hemoglobin (Exagamglogene autotemcel, Casgevy) or the production of modified hemoglobin A (HbA^T87Q^) (Lyfgenia) were FDA-approved based on encouraging results regarding their efficacy in reducing VOC ^16–19^. However, long term efficacy, progressive organ decline, and effects on life expectancy remain to be investigated. Moreover, patients awaiting or ineligible for gene therapy treatments will have continued need for other therapies.

Besides physical obstruction, lysis of stiff RBCs leads to the release of HbS and heme, which in turn provoke VOC and ACS ^20,21^. Once in the extracellular space, heme acts as a potent prothrombotic and proinflammatory damage-associated molecular pattern (DAMP) driving thrombosis and inflammation in different organs ^22^. Extracellular heme was shown to induce ACS in SCD mice and blocking P-selectin reduces lung injury ^21,23^. Platelet-neutrophil aggregates promote lipopolysaccharide (LPS)-induced VOC in the lung arterioles of SCD mice and blocking P-selectin resolves pulmonary arteriole microemboli ^12^, however thrombo-inflammation and embolic NETs arriving to the lungs persist in challenged SCD mice treated with P-selectin blocking antibody ^13^. Therefore, there is an urgent need to identify new targets to limit VOC and thrombo-inflammation and to improve ACS and organ perfusion in patients with SCD.

Free oxidized heme (hemin) was shown to induce neutrophil extracellular trap (NET) formation ^24,25^ and endothelial cell activation through toll-like receptor (TLR)-4^20^, and platelet activation through C-type lectin receptor (CLEC)-2 and glycoprotein (GP) VI ^26–28^, all together contributing to thrombo-inflammation in SCD. Many of the receptors involved in VOC and thrombo-inflammation in SCD are regulated by spleen tyrosine kinase (Syk). Syk is ubiquitously expressed across several cell types relevant to SCD pathophysiology, including hematopoietic lineages, endothelial cells, and epithelial cells. Syk was shown to regulate neutrophil adhesive functions ^29^, P-selectin-mediated NET formation and P-selectin glycoprotein ligand (PSGL)-1 activation ^30,31^, TLR4 signal transduction ^32^ and NLRP3 inflammasome activation ^33^, all of which described to contribute to VOC in SCD. Moreover Syk blockade reduced hemin-mediated platelet activation and aggregation *in vitro* ^27^ and decreased RBC adhesion and deformability in SCD blood *ex-vivo* ^34^. However, whether Syk inhibition reduces ACS and VOC in SCD mice is not known. In this study, we assessed whether blocking Syk in SCD mice alters thrombo-inflammation in the lungs, liver, spleen, and kidney of SCD at basal state and following hemin treatment and its effect on vascular integrity and organ perfusion.

## Methods

### Additional methods in supplementary file

#### Human blood

Venous blood was drawn from healthy, consenting, drug-free volunteers into 3.2 % trisodium citrate BD Vacutainers (Becton Dickinson, UK). Ethical approval was granted by the University of Birmingham Research Ethics Committee (ERN_11-0175).

#### Mice

Berkeley SCD mice ^35^ were purchased from Jackson Laboratories. Age-matched males and females sickle (Hba^0/0^ Hbb^0/0^ Tg(Hu-miniLCRα1^G^γ^A^γδβ^S^) and non-sickle control mice (Hba^0/0^ Hbb^0/+^ Tg(Hu-miniLCRα1^G^γ^A^γδβ^S^) (9-14 weeks) were used. All experiments were performed in accordance with UK law (Animal Scientific Procedures Act 1986) with approval of the local ethics committee and UK Home Office approval under PPL PP5212425 granted to the University of Birmingham.

#### Intravital imaging of hemin-induced thrombo-inflammation and perfusion in ventilated lungs

Experiments were performed under terminal anesthesia through intraperitoneal injection of ketamine hydrochloride (100 mg/kg) and medetomidine hydrochloride (10 mg/kg). Mice were intubated and ventilated with medical oxygen on a MiniVent rodent ventilator (stroke volume: 220 μL, respiratory rate: 130 breaths/min; Biochrom Ltd./Harvard Apparatus). VOC was induced by intravenous injection of 20µmol/kg of hemin (Frontiers specialty chemicals) via a retroorbital route. A concentrated solution of hemin was prepared in 50mM NaOH, diluted 10 times in PBS, pH adjusted to 7.3 and filtered using a 0.22µm filter before injection to mice. EDTA anticoagulated blood was collected via retroorbital route and cell count measured using an automated hematology analyzer, ABX Pentra 80 (HORIBA). In some experiments, Syk inhibitor BI-1002494 (4 or 20 mg/kg in 200µl saline containing a final concentration of 10% DMSO) (Boehringer Ingelheim, Germany) was injected intraperitoneally to mice 30 minutes before injection of hemin. Vehicle (saline-10%DMSO) was used as control. Real-time intravital observations were performed as previously described ^36,37^. Briefly, a thoracic window (steel stabilizer) was used to apply gentle negative pressure (∼20 mmHg) to a region of the surface of the left lung, allowing stabilization with maintained ventilation ^37^. To simultaneously image endogenous neutrophils and platelets, PE anti– mouse Gr-1 (Ly-6G/Ly-6C) (clone RB6-8C5, BioLegend) and Dylight649 anti-GPIb-beta (GPIbβ) (X-649, Emfret) were injected 5 min prior to imaging. Intravital imaging was performed using an upright microscope (BX61WI, Olympus) equipped with a Nipkow spinning disk confocal head (Yokogawa CSU) and an Evolve EMCCD camera (Photometrics) ^38^. The first 1-min capture was performed at baseline, followed by hemin injection (20 µmol/kg). Images were then captured every 15 minutes in the same area for 1 hour at both 10x and 40x magnification. At the end of the imaging period, the stabilizer was moved to the left kidney, spleen, and liver for additional images. Data were captured, stored, and analyzed using Slidebook 6 software (Intelligent Imaging Innovations). Neutrophils and platelet aggregates/microthrombi at 10x magnification were quantified by using segment masks on PE-Gr-1 and Dylight649 anti-GPIbβ staining, respectively. Integrated fluorescence density, accounting for size and fluorescence intensity, was calculated using ImageJ. Laser speckle contrast imaging (LSCI) was used to quantify left pulmonary blood flow in mice as previously described ^39^. LSCI and tissue histology are detailed in supplementary methods.

#### Human platelet aggregation

Human washed platelets were prepared and aggregation performed as previously described ^27^. Measurement and analysis of neutrophil adhesion and NETosis are detailed in supplementary methods.

#### Statistical analysis

All data are presented as mean±SD unless otherwise stated. The statistical significance between 2 groups was analyzed using student’s unpaired t-test and the statistical difference between multiple groups using one-way ANOVA with Tukey’s or Dunnett’s multiple comparisons test or the Kruskal-Wallis test as stated in the figure legends using Prism 8 (GraphPad Software Inc, USA).

## Results

### Extracellular hemin induced platelet and neutrophil recruitment to the pulmonary vasculature and impaired lung perfusion in SCD mice

Extracellular hemin was shown to induce VOC and ACS in different humanized SCD mice (Berkeley, Townes) ^20,21^. Hemin injections precipitate hemolytic and thrombo-inflammatory responses resembling sickle cell ACS in humanized SCD mice. A dose of 75 µmol/kg of hemin induced 100% lethality whereas a lower dose (17.5 µmol/kg) did not induce lethality within 2 hours of treatment in SCD mice ^21^. We therefore used a dose of 20 µmol/kg to track neutrophil and platelet recruitment to the lung using intravital imaging and to measure the perfusion using LSCI.

At baseline, platelet and RBC counts were lower in SCD mice compared to controls whilst white blood cell (WBC) and neutrophil counts were higher in SCD mice (Supplementary Figure 1A-D). Lung and kidney perfusion was not altered in SCD mice compared to control whereas liver and spleen perfusion was significantly reduced in SCD mice (Supplementary Figure 1E-H). As the number of SCD mice generated is limited and the aim of the study is to assess the effect of Syk inhibitor BI-1002494 on ACS in SCD mice, we assessed whether cell recruitment in mice treated with vehicle had altered responses compared to untreated mice. Our data confirmed that pretreatment with vehicle did not alter platelet or neutrophil recruitment to the lung microcirculation when comparing vehicle-treated to untreated control and SCD mice (Supplementary Figure 2A-D). Vehicle treatment was subsequently used as the control in this study. Intravital imaging of the lungs showed higher basal neutrophil adhesion in the microcirculation in SCD mice compared to controls (Figure 1A-D). Despite increased neutrophil and platelet recruitment in SCD mice at baseline, lung perfusion in these mice was not altered compared to controls (Supplementary Figure 1E). Hemin triggered progressive neutrophil recruitment to the pulmonary microcirculation in control mice (Figure 1A-E). Neutrophil recruitment, which was significantly higher in SCD mice at baseline, was further increased following hemin treatment. Overall neutrophil recruitment to the lung over 1h of imaging was higher in SCD compared to control mice, resulting in a high coverage of the pulmonary microvasculature (Figure 1E). Injection of hemin increased unstable platelet recruitment (Figure 1A, B) and the formation of small aggregates in control mice (Figure 1A-H). Stable platelet aggregates and large platelet emboli were also observed in pulmonary microcirculation of hemin-treated SCD mice (Figure 1A, B, Supplementary video 1 and 2). In control mice treated with hemin, platelets adhered and embolized without apparent stable binding. Large platelet emboli were detected in SCD mice along stable adherent platelets (Figure 1A-B, Supplementary video 1 and 2). Injection of hemin in control mice did not alter lung perfusion compared to baseline as measured using LSCI but reduced lung perfusion in SCD mice (Figure 1I, J). Increased platelet and neutrophil adhesion to the pulmonary microcirculation was associated with decreased perfusion following hemin treatment in SCD mice. In conclusion, hemin exacerbated platelet and neutrophil recruitment to the lung microcirculation and altered lung perfusion.

**Figure 1:**
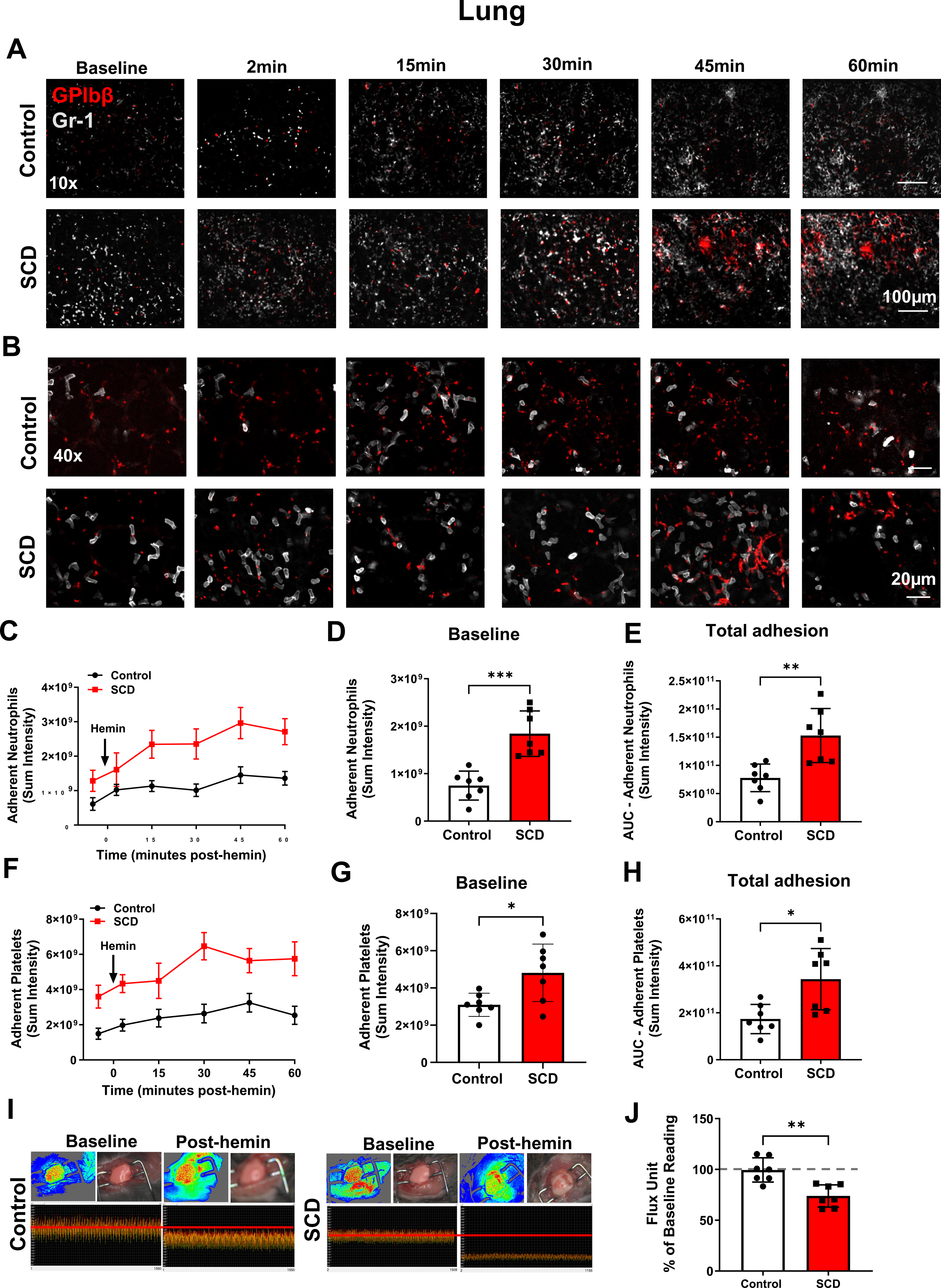
Extracellular hemin increased neutrophil recruitment and stable platelet aggregates in the pulmonary microcirculation. Hemin (20µmol/kg, I.V.) was injected in control and SCD mice. Representative intravital images of the breathing lungs showing adherent neutrophils (Gr-1; grey) and platelets (GPIbβ; red) in the pulmonary microvasculature over a time course of 60 min at (A) 10x and (B) 40x magnification. (C) Quantitative analysis of the intravital data for adherent neutrophils over the time course. (D) Adherent neutrophil at baseline measured as sum intensity and (E) and the area under the curve (AUC) analysis over a time course of 60 min. (F) Quantitative analysis of the intravital data for platelets over the time course. (G) Quantitative analysis of platelet sum intensity at baseline and (H) AUC analysis for platelet recruitment over a time course of 60 min. (I) Representative LSCI images showing flux data, flux heat maps and a corresponding photo of the breathing lungs. (J) Quantitative analysis of flux unit readings as a percentage of baseline values obtained by LSCI. The statistical significance between 2 groups was analyzed using an unpaired t-test. *p < 0.05, **p < 0.01, ****p<0.0001.

### Organ-specific alteration in neutrophil and platelet recruitment following hemin injection

We next assessed whether hemin induced a systemic alteration in cell recruitment or alteration in organ perfusion. We compared platelet and neutrophil recruitment at baseline and 1h post-hemin injection in the kidney, liver and spleen using intravital microscopy. At baseline, neutrophil and platelet retention were higher in the kidney of SCD compared to control (Figure 2A-C). Injection of hemin in control mice did not alter platelet or neutrophil recruitment to the kidney while hemin increased neutrophil recruitment to the kidney of SCD mice (Figure 2A-C). Neutrophil and platelet presence in the spleen was comparable at baseline between control and SCD mice (Figure 2D-F). Following hemin injection, splenic neutrophil and platelet numbers were unaltered in control mice whereas hemin-induced reductions in neutrophils and platelets were detected in SCD mice, potentially reflecting release of platelets and neutrophils stored in the spleen (Figure 2D-F). Imaging of the liver microcirculation revealed reduced neutrophils presence in SCD mice at baseline compared to control (Figure 2G-I). Hemin injection did not alter neutrophil recruitment in control mice but reduced neutrophils in SCD mice. Conversely, hemin decreased platelets in the liver of control but not SCD mice (Figure 2G-I).

**Figure 2:**
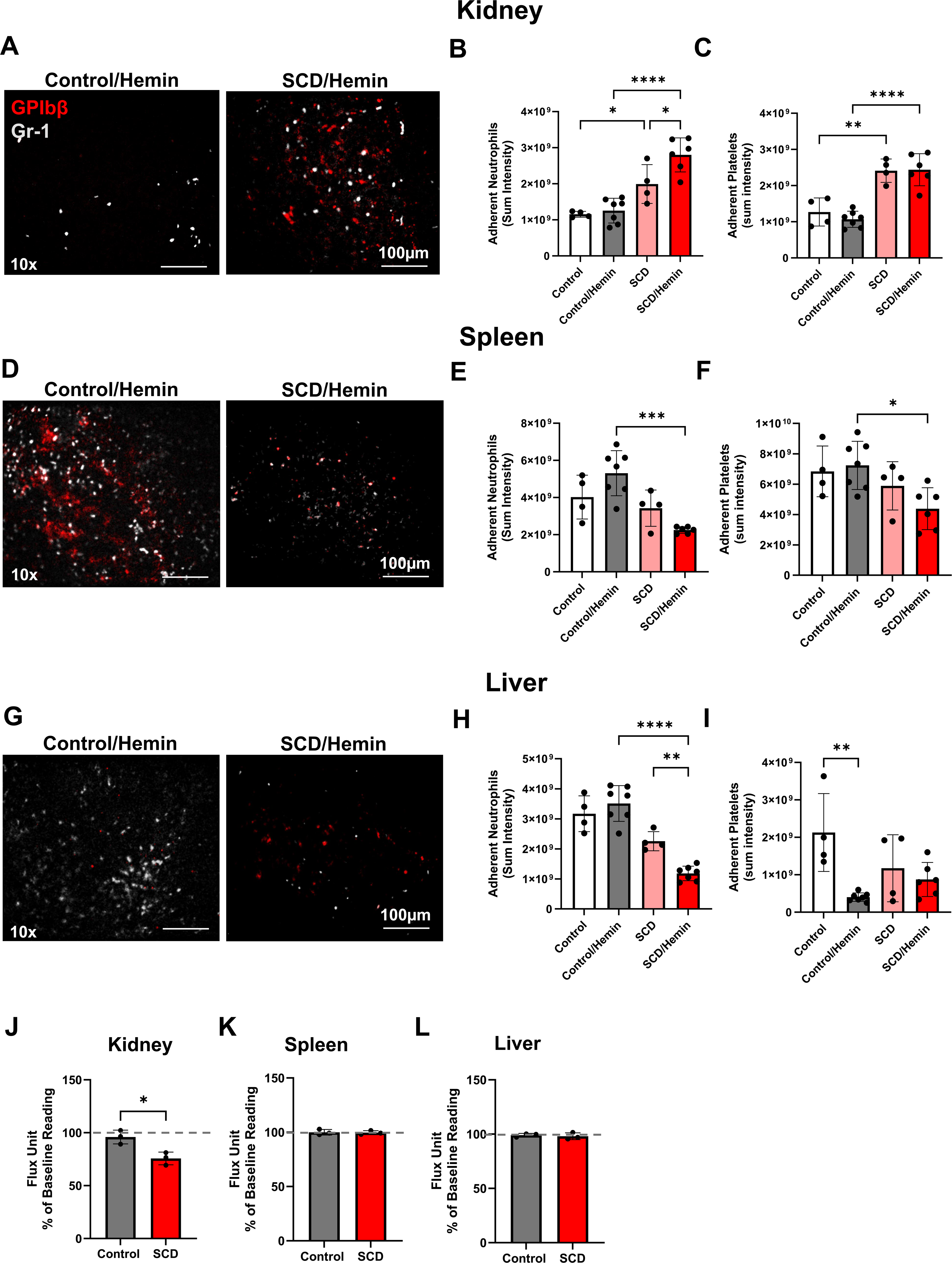
Extracellular hemin increased neutrophil recruitment to the kidney in SCD but impaired their recruitment to the spleen and liver. Hemin (20µmol/kg, I.V.) was injected into control and SCD mice. Neutrophil and platelet recruitment to the kidney, spleen and liver was analyzed at 60min post-hemin and imaged at 10x magnification. (A) Representative intravital images of the kidney showing adherent neutrophils (Gr-1; grey) and platelets (GPIbβ; red) in the renal microcirculation. Quantitative analysis of the intravital data for (B) total adherent neutrophils and (C) platelets before and 1h following hemin injection. (D) Representative intravital images of the splenic microcirculation showing adherent neutrophils (Gr-1; grey) and platelets (GPIbβ; red) in the renal microcirculation. Quantitative analysis of the intravital data for (E) total adherent neutrophils and (F) platelets before and 1h following hemin injection. (G) Representative intravital images of the liver showing adherent neutrophils (Gr-1; grey) and platelets (GPIbβ; red) in the hepatic microcirculation. Quantitative analysis of the intravital data for (H) total adherent neutrophils and (I) platelets before and 1h following hemin injection. Quantitative analysis of flux unit readings as a percentage of baseline values obtained by LSCI for the (J) kidney, (K) spleen and (L) liver. The statistical significance between 2 groups was analyzed using an unpaired t-test and the difference between multiple groups analyzed using ordinary one-way ANOVA with Tukey’s multiple comparison test. *p < 0.05, **p < 0.01, ****p<0.0001.

We next assessed organ perfusion at baseline and following hemin treatment in control and SCD mice. Injection of hemin reduced kidney perfusion in SCD but not in control mice (Figure 2J). No alteration in liver or spleen perfusion were observed in control or SCD mice compared to control, with both organs showing lower perfusion in SCD at baseline compared to control (Figure 2K-L, Supplementary Figure 1G-H). Therefore, hemin impaired lung and kidney perfusion is associated with increased neutrophil and platelet recruitment to these organs. These data show that organ-specific alterations in neutrophil and platelet recruitment occur in SCD mice associated with impaired perfusion.

### Hemin-induced bleeding and thrombo-inflammation in the ventilated lungs is independent of local RBC sickling and adhesion

Hemin injection in sickle mice was shown to induce vascular congestion, edema, alveolar wall thickening and hemorrhage in the lungs ^21^. In our study, histological analysis of the formalin-fixed and paraffin-embedded sections from ventilated lungs and other organs post-intravital imaging showed organ-specific RBC sickling. Iron deposition was observed in the lungs of unchallenged SCD mice (brown spots) (Figure 3A, magnification). Hemin increased vascular congestion in control mice which was exacerbated in SCD mice. Moreover, hemin induced bleeding in the lungs of SCD mice without evidence of RBC sickling, possibly due to lung ventilation and local delivery of oxygen to the pulmonary microcirculation (Figure 3A-B). Conversely, hemin induced RBC adhesion in the livers of control mice were exacerbated in SCD mice. RBC sickling and adhesion was significantly higher in the liver of SCD mice, both in the sinusoids and large vessels (Figure 3C). Sickled and adherent RBC were also observed in kidneys of ventilated SCD mice treated with hemin (Figure 3D). Sickled RBC were also observed in the heart (not shown). Significant alteration in the splenic white and red pulp of SCD mice were observed with increased RBC and trapping observed following hemin injection (Figure 3E). These data show that hemin induced bleeding in the lung of ventilated mice, as well as vascular congestion without increasing RBC sickling and adhesion, suggesting that the impairment in perfusion is likely related to neutrophil adhesion, which can lead to inflammatory bleeding.

**Figure 3:**
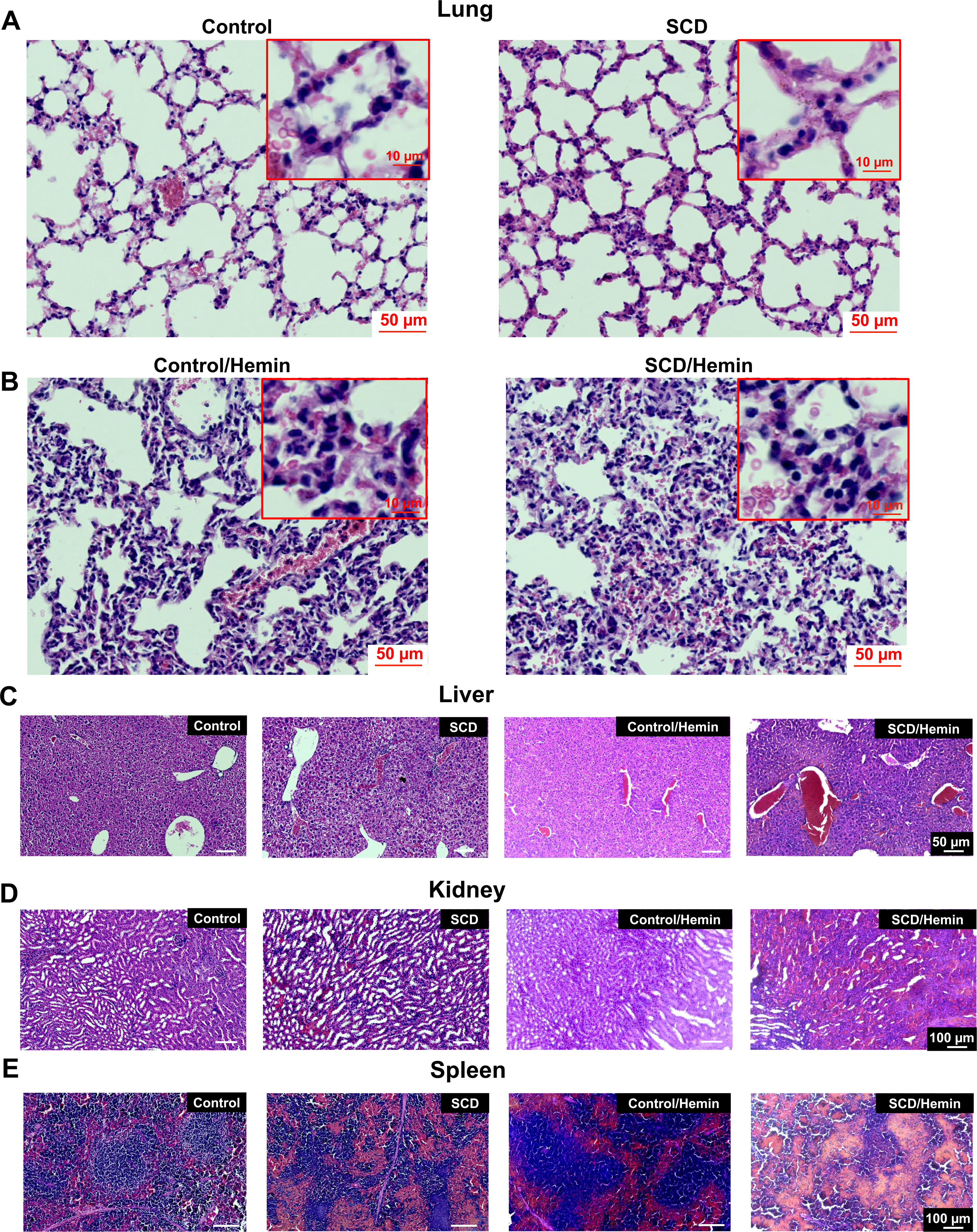
Extracellular hemin injection in ventilated mice induced bleeding in the lungs and induced RBC sickling and VOC in the liver and kidney. Organs from unchallenged control or SCD mice were used for baseline tissue analysis. Organs were collected from ventilated mice 1h following hemin injection and processed for H&E staining. (A, B) Lung, (C) liver, (D) kidney, and (E) spleen whole sections were imaged using Zeiss Axioscan microscope and analyzed using Zen software (n=3).

### Syk inhibitor BI-1002494 inhibited hemin-induced human platelet aggregation and hemin-induced neutrophil adhesion

Neutrophil adhesion to fibrinogen ^29^, NET formation by selective stimuli such as uric acid ^40^ and hemin-mediated platelet activation ^27^ are regulated by Syk. NETs were detected in the lungs of SCD mice and hemin-induced NETs were shown to contribute to the pathogenesis of SCD ^24^. Moreover, neutrophils were shown to trigger bleeding at the site of inflammation ^41^, a mechanism which could contribute to bleeding in the lung of SCD mice. We therefore assessed the effect of Syk inhibition on hemin-induced thrombo-inflammation in SCD mice. We first tested the effect of two selective Syk inhibitors, BI-1002494 and PRT-060318, on human neutrophil adhesion on VWF and fibrinogen *in vitro,* hemin-induced human platelet aggregation and NET formation. Neutrophil adhesion to VWF and fibrinogen was significantly increased by addition of hemin (5µM) but not by TNF-α (Figure 4A, B). BI-1002494 and PRT-060318 inhibited neutrophil adhesion to VWF or fibrinogen, in the presence or absence of TNFα (not shown) or hemin (Figure 4C, D). Hemin (5 µM) did not induce NET formation in healthy neutrophils while addition of platelets in the presence of hemin significantly increased NET formation (Figure 4E, H). We further focused on BI-1002494 to validate its efficacy on blocking NETosis. NETosis driven by activated platelets was inhibited by pretreatment of platelets with BI-1002494. PMA-induced NETosis, a Syk-independent mechanism, was not affected by BI-1002429 while uric acid-induced NETosis, a Syk-dependent mechanism, was significantly inhibited (Figure 4E-G). In line with our previous study ^27^, both inhibitors (10µM) abolished hemin-induced platelet aggregation as assessed using light transmission aggregometry (Figure 4I). Therefore, this data suggests that Syk inhibition blocks neutrophil adhesion and platelet activation triggered by hemin and limits Syk-dependent NETosis.

**Figure 4:**
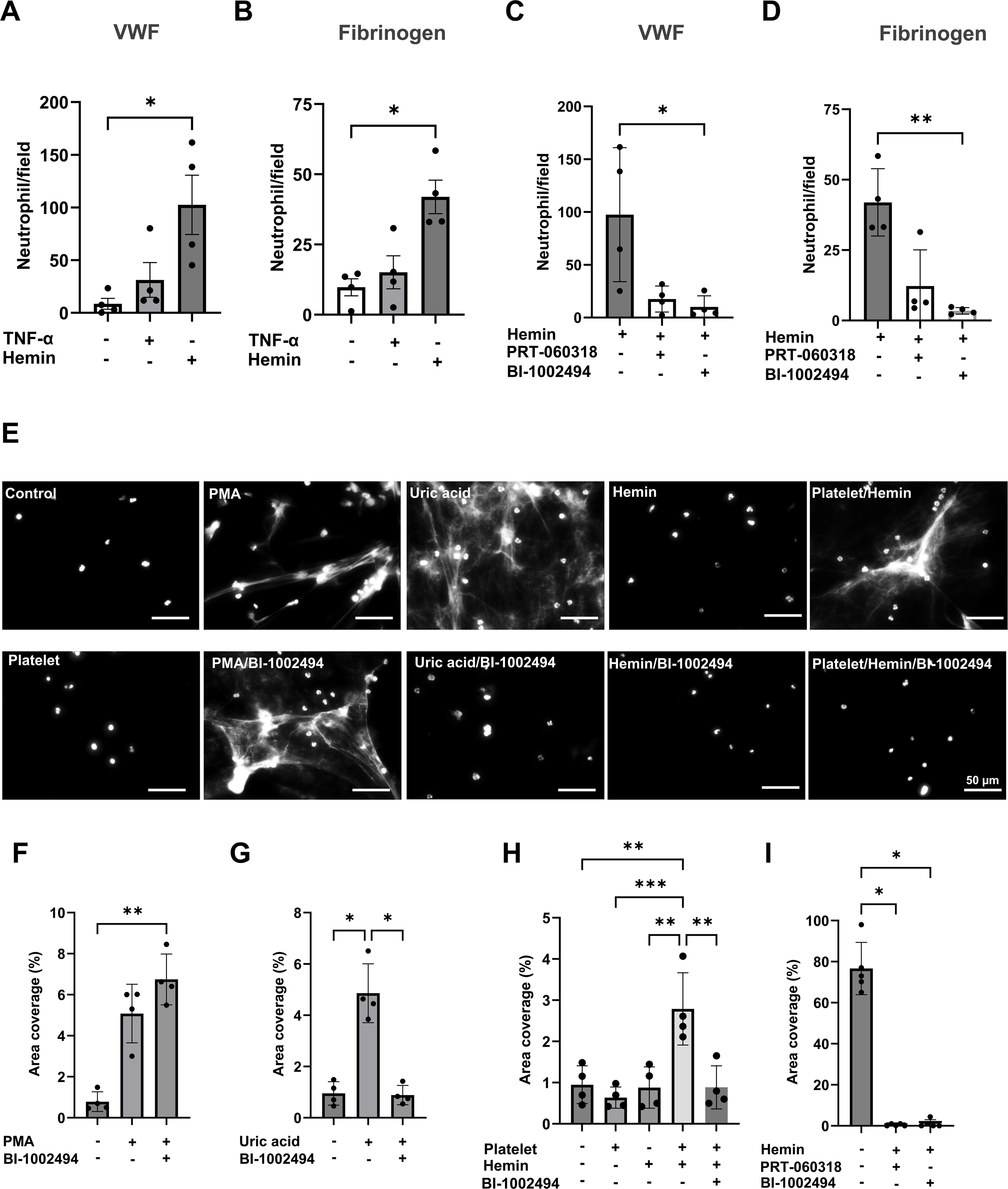
Syk inhibitors BI-1002494 and PRT-060318 reduced neutrophil adhesion, NETosis and platelet aggregation by hemin. (A, B) Purified neutrophil adhesion (30min at 37 **°**C) on (A) immobilized VWF (100 µg/ml) or (B) fibrinogen (200µg/ml) was assessed in the presence or absence of TNF-α (10ng/ml) or hemin (5µM) (n=4 independent experiments, average of 5 images). (C, D) Neutrophils were pretreated with Syk inhibitors, PRT-060318 or BI-1002494 (10µM), prior to incubation with hemin on (C) VWF or (D) fibrinogen. (E-H) Neutrophil extracellular trap formation *in vitro*. (E) Representative image of DNA stain using Sytox Green. Neutrophils were treated with PMA (100nM), uric acid (20µg/ml), non-activated or hemin-activated platelets for 3h. BI-1002494 was used at 10 µM. (F-H) Area coverage of DNA was quantified and averaged from 5 separate fields per condition in each donor and the results shown as mean±SD of 3-4 independent donors and experiments. (I) Human washed platelet aggregation by hemin (5µM) was assessed by light transmission aggregometry. Platelets were pretreated for 10min at 37 **°**C with PRT-060318 (10µM) prior to stimulation with hemin. The significant difference between multiple groups was assessed using Kruskal-Wallis multiple comparison test compared to control. *p < 0.05, **p < 0.01, ****p<0.0001.

### Syk inhibitor BI-1002494 inhibited platelet and neutrophil recruitment to the lung and improved lung perfusion

Previously, prophylactic or therapeutic administration of BI-1002494 (100 mg/kg) by oral gavage was shown to protect mice against arterial thrombosis and reduced brain infarct in an experimental model of ischemic stroke without altering hemostasis ^42^. In the current study, two doses of BI-1002494, 20 mg/kg or 4mg/kg, were administered via intraperitoneal (IP) injection 30 min prior to hemin administration. The assessment of the efficacy of two doses was related to potential side effects of high dose Syk inhibition in SCD mice due to inflammatory bleeding. BI-1002494 treatment did not alter basal lung perfusion in SCD mice (Supplementary Figure 3A). BI-1002494, at 20mg/kg or 4mg/kg, inhibited baseline and hemin-induced platelet and neutrophil recruitment to the lung and improved perfusion following hemin administration in SCD mice (Figure 5A-G). Moreover, BI-1002494 treatment restored lung perfusion in hemin-treated mice (Figure 5H). Therefore, we observed that low-dose BI-1002494 (4mg/kg) is sufficient to inhibit neutrophil and platelet recruitment and to restore lung perfusion in SCD following hemin treatment.

**Figure 5:**
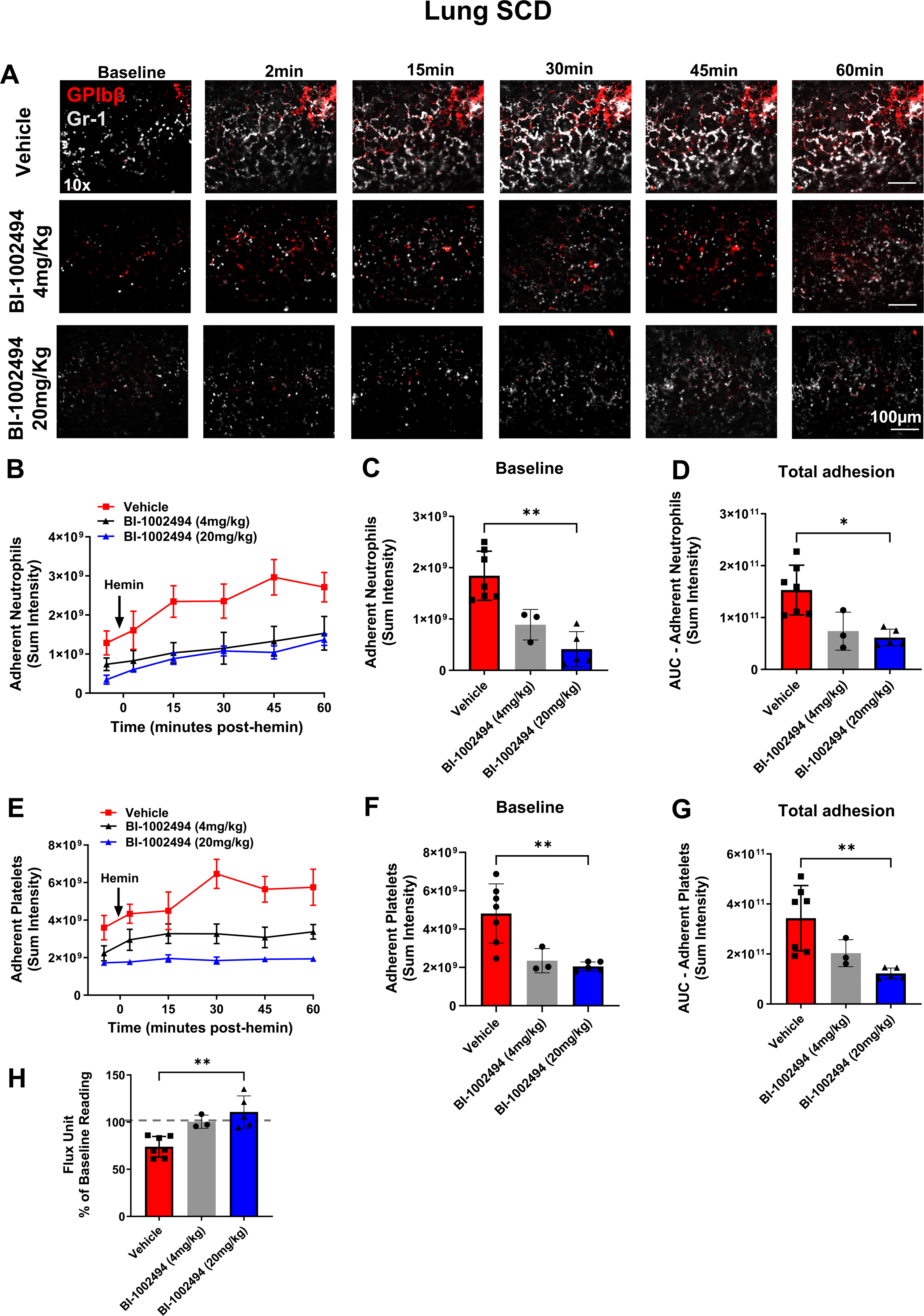
Syk inhibitor BI-1002494 inhibited neutrophil and platelet recruitment to the lungs in SCD at baseline and following hemin injection. BI-1002494 (4 or 20mg/kg) or vehicle were injected in SCD mice via intraperitoneal route 30 min before the injection of hemin (20µmol/kg, I.V.). (A) Representative intravital images of the ventilated lungs showing adherent neutrophils (Gr-1; grey) and platelets (GPIbβ; red) in the pulmonary vasculature over a time course of 60 min at 10x magnification. Quantitative analysis of the intravital data for adherent neutrophils (B) over the time course, at (C) baseline, (D) and the area under the curve (AUC) analysis over a time course of 60 min. Quantitative analysis of the intravital data for platelets (E) over the time course, at (F) baseline, (G) and AUC analysis over a time course of 60 min. (H) Quantitative analysis of flux unit readings as a percentage of baseline values obtained by LSCI. The statistical significance between multiple groups analyzed using Kruskal-Wallis test with multiple comparison compared to control. *p < 0.05, **p < 0.01, ****p<0.0001.

### BI-1002494 reduced platelet and neutrophil recruitment to the kidney and improved organ perfusion

We next assessed the effect of BI-1002494 on platelet and neutrophil recruitment and perfusion in the kidney, spleen, and liver following hemin treatment. At baseline, BI-1002494 did not alter the perfusion in the kidney, spleen or liver compared to vehicle-treated mice (Supplementary Figure 3B-D). Intravital imaging showed a decrease in platelet and neutrophil recruitment to the kidney in BI-1002494-treated SCD mice compared to vehicle-treated mice, independent of the dose tested (Figure 6A-C). Kidney perfusion was assessed in mice treated with 20mg/kg BI-1002494. LSCI results showed that the pretreatment of mice with BI-1002494 improved kidney perfusion following hemin treatment (Figure 6D). Conversely, a dose-dependent effect on platelet and neutrophil recruitment was observed in the liver. The impairment in neutrophil recruitment to the liver observed in SCD mice was reversed by 20mg/kg but not 4mg/kg of BI-1002494 (Figure 6E-G). At the higher dose, platelet recruitment to the liver was significantly increased in BI-1002494-treated mice compared to vehicle mice. BI-1002494 did not alter liver perfusion in SCD mice compared to vehicle-treated mice (Figure 6H). Finally, BI-1002494 (20mg/kg) reduced neutrophil recruitment to the spleen without altering platelet recruitment, while the dose of 4mg/kg had no effect on platelet or neutrophil recruitment (Supplementary Figure 4A-D). Therefore, targeting Syk improves hemin-induced thrombo-inflammation in the lung and kidney and the perfusion in these organs.

**Figure 6:**
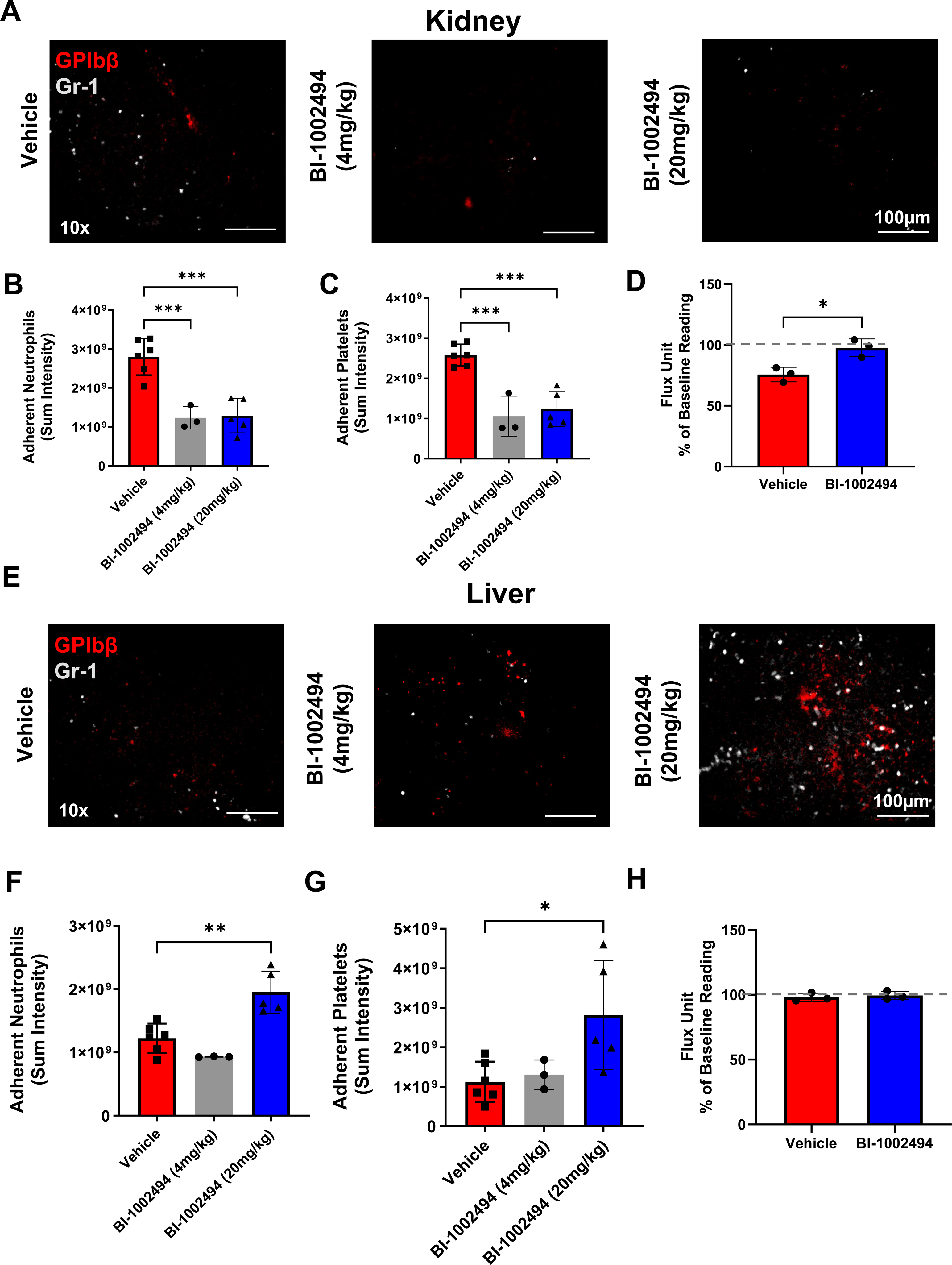
Syk inhibitor BI-1002494 reduced neutrophil and platelet recruitment to the kidney but increased their recruitment to the liver of SCD mice. BI-1002494 (4 or 20mg/kg) or vehicle were injected in SCD mice via intraperitoneal route 30 min before the injection of hemin (20µmol/kg, I.V.). Neutrophil and platelet recruitment to the kidney and liver was analyzed at 60min post-hemin and imaged at 10x magnification. (A) Representative intravital images of the kidney showing adherent neutrophils (Gr-1; grey) and platelets (GPIbβ; red) in the renal microcirculation. Quantitative analysis of the intravital data for (B) total adherent neutrophils and (C) platelets. (D) Quantitative analysis of flux unit readings as a percentage of baseline values obtained by LSCI for the kidney. (E) Representative intravital images of the liver showing adherent neutrophils (Gr-1; grey) and platelets (GPIbβ; red) in the hepatic microcirculation. Quantitative analysis of the intravital data for (F) total adherent neutrophils and (G) platelets at 60min post-hemin. (H) Quantitative analysis of flux unit readings as a percentage of baseline values obtained by LSCI for the liver. The statistical significance between 2 groups was analyzed using an unpaired t-test and the statistical significance between multiple groups analyzed using one-way ANOVA with Dunnett’s multiple comparison compared to control *p < 0.05, **p < 0.01, ****p<0.0001.

### Low concentrations of BI-1002494 inhibited hemin-induced inflammatory bleeding in the lung and reduced VOC in the kidney of SCD mice

Administration of hemin induced bleeding in the lungs of SCD mice. Histological analysis using H&E staining revealed that the lower concentration of BI-1002494 (4mg/kg) limited bleeding in the lungs of hemin-treated SCD mice while higher concentrations increased bleeding (Figure 7A). BI-1002494, at both concentrations, reduced RBC adhesion in the kidney (Figure 7B). No alteration of VOC in the liver was observed in the liver or spleen (Figure 7C, D). This data show that BI-1002494 (4mg/kg) is effective in reducing platelet and neutrophil recruitment to the lung, improving perfusion, and reducing bleeding risk associated with hemin injection in SCD mice.

**Figure 7:**
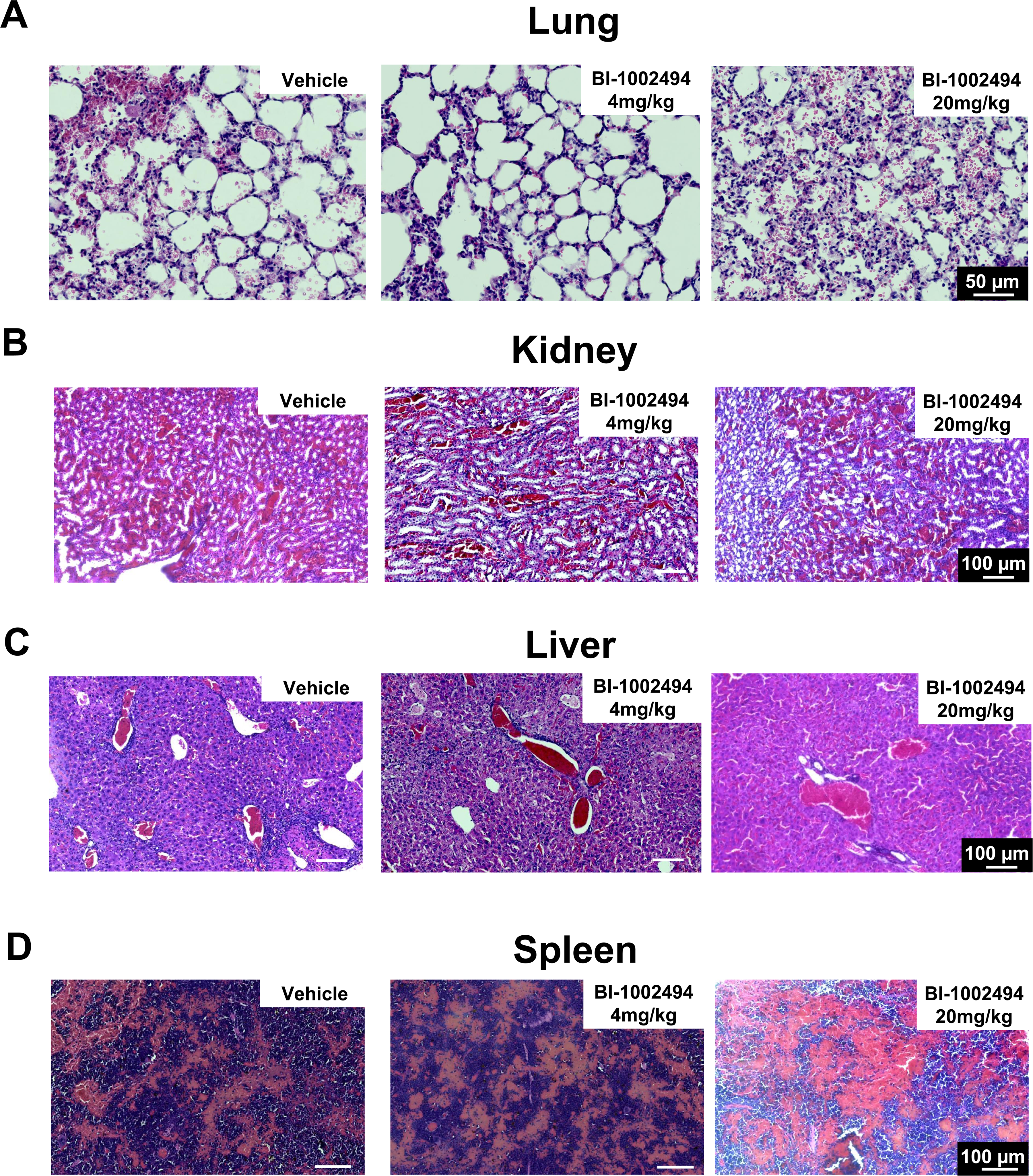
Low dose BI-1002494 reduced hemin-induced bleeding in the lung and decreased VOC in the kidney. BI-1002494 (4 or 20mg/kg) or vehicle were injected in ventilated SCD mice via intraperitoneal route 30 min before the injection of hemin (20µmol/kg, I.V.). Lung, kidney, liver and spleen were fixed 60 min post-hemin injection. H&E staining of (A) lung, (B) kidney, (C) liver and (D) spleen. Bar 100µm (n=3).

## Discussion

In this study, we investigated the effect of hemin on neutrophil and platelet recruitment in the lung, kidney, liver, and spleen of control and SCD mice using intravital imaging. We observed organ-specific effects of hemin on neutrophil and platelet recruitment as well as impairment of vascular integrity in the lung of SCD mice. We show that hemin promoted neutrophil and platelet recruitment to the lung of ventilated SCD mice and impaired perfusion. This was associated with bleeding and vascular congestion without RBC sickling and adhesion in the lung under these conditions. Hemin induced RBC sickling in other organs including the kidney, liver and spleen and resulting in their adhesion and red clot formation. The inhibition of Syk using BI-1002494 reduced neutrophil and platelet adhesion, reduced bleeding in the lung and improved perfusion. Increasing BI-1002404 concentration reduced thrombo-inflammation but increased bleeding in the lungs. However, the inhibition of Syk did not improve RBC sickling and VOC in the liver and other organs.

Human clinical trials exploring anti-inflammatory and anti-thrombotic therapies have thus far demonstrated limited efficacy in reducing morbidity in patients with sickle cell disease. While platelet and neutrophil contributions to VOC has been shown in multiple studies, blocking ADP receptor P2Y_12_ using prasugrel did not improve VOC in patients with SCD as shown by the DOVE trial ^43^. Despite encouraging early results ^44,45^, anti-P-selectin antibody Crizanlizumab did not outperform the placebo in the phase III randomized-controlled clinical trial STAND and was revoked by the European Medicines Agency ^45–48^. The limited efficacy of these therapies in reducing VOC in patients remains poorly understood. It is likely that multiple mechanisms support VOC and thrombo-inflammation with some degree of organ specificity and these mechanisms could be further modulated by the insults triggering VOC in patients. Indeed, by imaging different organs, we were able to observe organ-specific recruitment of neutrophils and platelets in response to hemin injection and that targeting Syk resulted in organ-specific alternations in thrombo-inflammation and perfusion.

Both free heme and RBC-derived microparticles loaded with heme were shown to activate endothelial cells and to support VOC in SCD ^49^. Characterization of the molecular mechanisms indicated that hemin-dependent endothelial cell activation is TLR4-dependent and blocking TLR4 signaling inhibited endothelial activation and mitigated VOC and ACS in SCD mice ^20,21^. Heme can directly activate platelets via immunoreceptor tyrosine-based activation motif (ITAM) signaling receptors CLEC-2 and GPVI ^27,28^. In a model of SCD, injection of hemin resulted in its accumulation in the kidney and triggered acute kidney injury whereas injection of hemin in control mice led to the accumulation of hemin in the liver ^50^. This could explain the increase in neutrophil recruitment in the liver in control but not in SCD mice in our model, as well the increased neutrophil and platelet recruitment to the kidney of SCD mice. The increase in neutrophil recruitment to the lung and kidney could be related to increased susceptibility of specific vascular beds to hemolysis and inflammatory stimuli. Indeed, SCD mice have higher levels of vascular cell adhesion protein-1 (VCAM-1) and intercellular adhesion molecule-1 (ICAM-1) which are increased by 3-5-fold in the lungs, indicating a high level of inflammation and endothelial cell activation in the lung of SCD mice ^51^. Increased endothelial activation can increase neutrophil and platelet recruitment, independent of RBC sickling and adhesion. Platelet-neutrophil complexes and embolization were shown to mediate VOC in the lungs of SCD mice following LPS injection ^12^. This organ-specific effect could also be related to the increased susceptibility of specific endothelial beds to heme, as shown with glomerular endothelial cells ^52^. These data support organ-dependent mechanisms of platelet and neutrophil recruitment in SCD mice and support the need to target selective pathways involved in different vascular beds, which can lead to organ-specific mechanisms of VOC and ACS.

Various lines of evidence support the hypothesis that Syk inhibition in SCD mice has the potential to reduce thrombo-inflammation, RBC sickling and VOC. The Syk inhibitor Fostamatinib is currently used clinically to treat immune thrombocytopenia (ITP) and short and long-term targeting Syk was shown to be safe in patients with ITP ^53,54^. In the context of SCD, targeting of Syk can potentially inhibit multiple key pathways involved in VOC and organ damage. First, activation of ITAM receptors by heme/hemin in platelets leads to Syk phosphorylation, PLCƔ2 activation and platelet activation ^55^. Second, Syk in neutrophils is required for their adhesion ^29^, and their recruitment to the lung vasculature in VOC in SCD mice ^12^. Third, engagement of PSGL-1 by P-selectin and E-selectin transactivates Syk in leukocytes and reduces neutrophil rolling velocities on E-selectin/ICAM-1 ^30,56–59^. Moreover, neutrophil interaction with immobilized P-selectin triggers Syk activation ^60^. Fourth, Syk phosphorylation was also found to be elevated in deoxygenated sickle RBCs and Syk inhibition, was shown to block Band 3 tyrosine phosphorylation preventing the release of extracellular vesicles, inhibiting RBC adhesion to heme-activated endothelial cells and increasing SCD RBC deformability in *ex-vivo* flow models ^34^. Fifth, heme can induce NLRP3 inflammasome activation and IL-1β secretion, a mechanism dependent on Syk, mitochondrial ROS and potassium influx ^61^. Furthermore, targeting Syk mitigates heme-provoked platelet activation and aggregation, underscoring its potential utility for dampening multiple facets of SCD vasculopathy ^27^. Intravital imaging in different organs provided new global information for Syk-dependent mechanisms driving neutrophil and platelet recruitment to different organs in response to hemin. Our data shows that impaired lung perfusion following hemin treatment is not regulated by RBC sickling but rather by neutrophil and platelet recruitment. Syk inhibition decreased thrombo-inflammation in the lungs and kidneys but did not alter RBC sickling and VOC in the liver. The development of organ-specific mechanisms might represent a limiting factor in using a single drug to limit thrombo-inflammation and reduce RBC sickling. As several oral Syk inhibitors are already undergoing evaluation in clinical cancer trials, and the Syk inhibitor Fostamatinib is approved for the treatment of patients with ITP, the efficacy of Syk inhibition in reducing thrombo-inflammation in the lung and kidney might open new possibilities to target VOC in SCD.

Our study has some limitations regarding the inhibitor used, the dose and route of administration. Future studies aiming to test different doses of Syk inhibitors, which are clinically approved such as Fostamatinib or currently evaluated in clinical trials, will provide critical information on the optimal dose of inhibitor to use to decrease thrombo-inflammation without impairing vascular integrity. The bleeding side effect of higher doses of Syk inhibitor used in this study is still not understood, but could be related to the route of administration and increased susceptibility of SCD mice to Syk inhibition or off-target effect on other kinases. Platelets maintain vascular integrity in the lungs following inflammatory challenges, however the contribution of platelet receptors to inflammatory hemostasis is stimulus dependent ^55,62^. CLEC-2 and GPVI deficiency did not alter vascular integrity in the lung of mice treated with LPS ^63^, however GPVI limits inflammatory bleeding following *Klebsiella pneumonia* infection ^64^. Platelet recruitment to the lung following LPS injection in wild-type mice is independent of neutrophils, PSGL-1 or P-selectin ^65^. The effect of different doses of BI-1002494 and other Syk inhibitors on vascular integrity in response to hemin and possibly other triggers of VOC requires further investigation. Moreover, the efficacy of long-term treatment of SCD mice with Syk inhibitors on progressive organ decline remains unknown. Finally, as multiple triggers and interventions were shown to induce VOC in SCD, including hemin, free hemoglobin S, LPS, hypoxia/oxygenation, cold and stress, future work assessing the efficacy of Syk inhibition under with different triggers of VOC would allow to assess its efficacy under different conditions.

In conclusion, this study is a proof-of-concept suggesting that of Syk inhibition could be beneficial in reducing thrombo-inflammation and ACS in SCD. It is possible that using low dose Syk inhibitors in conjunction with drugs decreasing RBC sickling will provide a better protection to reduce VOC and thrombo-inflammation and improve organ perfusion and function.

## Acknowledgments

J.R. holds a British Heart Foundation (BHF) Intermediate Fellowship (FS/IBSRF/20/25039). This research was partially supported by a BHF project grant for J.R (PG/21/10737). A.B holds a BHF senior Fellowship (FS/19/30/34173). S.P.W. holds a BHF Chair (CH03/003). JEA was supported by a BHF Project grant awarded to NK (PG/21/10574). The National Institute of Health and Care Research (NIHR) Birmingham Biomedical Research Centre (NIHR203326) and the BHF Accelerator (AA/18/2/34218) have supported the University of Birmingham Institute of Cardiovascular Sciences where this research is based. The opinions expressed in this paper are those of the authors and do not represent any of the listed organizations.

## Authorship Contribution

JEA performed intravital and laser speckle experiments, collected, analyzed data and contributed to data interpretation; GP, NM, VT, BG performed experiments and analyzed data; SJC developed methodology for intravital imaging; SPW, AB, JDD, PLRN contributed to experimental design and data interpretation; DK and NK provided key technical expertise and support for intravital imaging and LSCI and analysis, J.R. designed research and experiments, analyzed data, prepared figures and wrote the manuscript; all authors read and approved the paper.

