## Supplementary file for "Spleen tyrosine kinase inhibition mitigates hemin-induced thromboinflammation in the lung and kidney of sickle cell mice"

### Supplementary methods and figures:

#### Laser speckle contrast imaging of the ventilated mouse lungs

Laser speckle contrast imaging (LSCI) quantified left pulmonary blood flow in mice as previously described [31]. Following the surgical preparation described earlier, the LSCI device (moorFLPI-2; Moor Instruments, UK) was positioned above the exposed left lung. A defined area for flux data collection was marked at baseline and 1-hour post-hemin treatment. Additional recordings of the left kidney, spleen, and liver were taken both at baseline and 1-hour post-hemin treatment. Using the manufacturer-supplied image software (mFLPI2Measure V2.0; mFLPIReview V5.0), 1,000 frames were captured at each time point with a frame rate of 25 Hz and spatial processing (sliding window, time constant: 0.1 s). In-house-developed Basic Speckle Analysis software (SpAn; open source, available online at https://github.com/kavanagh21/SpAN) facilitated the identification and compilation of flux values.

#### Histology

Lung, kidney, spleen, and liver were collected from unchallenged control and SCD mice and from mice following intravital imaging procedures. Organs were immediately fixed in paraformaldehyde (PFA 4%) overnight, dehydrated and embedded in paraffin. Formalin-fixed paraffin-embedded (FFPE) sections (6µm) were processed for H&E staining (Abcam). Images were analyzed using Zeiss Axio Scan.Z1 microscope and ZEN software.

#### Human platelet aggregation

Human washed platelets were prepared and aggregation performed as previously described [25]. Platelets ($2x10^8$/ml) were preincubated with BI-1002494 (Boehringer Ingelheim) (10µM) and PRT-060318 (10µM) were incubated for 15 minutes at 37 °C prior to stimulation with hemin (5µM). Platelet aggregation assessed in the presence of 2.5 mM $CaCl_2$ for 6 minutes using PAP-8E aggregometer.

**Neutrophil adhesion and NETosis**

Neutrophils were isolated from EDTA-anticoagulated blood using histopaque 1077 and 1119 (Sigma-Aldrich). For neutrophil adhesion, fibrinogen (200 µg/ml) and VWF (100 µg/ml Wilfactin, LFB, France) were coated on 24-well plate for 1h at 37 °C, blocked with bovine serum albumin (BSA) fraction V (7.5%) for 1 hour and washed with PBS before addition of purified neutrophils (0.5 million/well) for 30 minutes at 37 °C. TNF-α (10ng/ml, Peprotech), hemin (5µM) were added on neutrophils and cells incubated for 30 minutes at 37 °C. In some conditions, neutrophils were preincubated with BI-1002494 or PRT-060318 (10µM) for 15min at 37 °C prior to addition to TNF-α and hemin. After 30 minutes incubation, cells were washed with PBS and neutrophil adhesion assessed using EVOS microscopy. The average neutrophil count from 5 representative images from each donor was analyzed.

For NETosis experiments, neutrophils (200,000/well) were added on poly-L-Lysine for 30 minutes before stimulation with phorbol myristate acetate PMA (100nM) or uric acid (20 µg/ml) for 3 hours. For experiments in the presence of platelets, human washed platelets (20 million) were stimulated with hemin (5µM) for 20 minutes before addition on neutrophils. Unstimulated platelets were used as control. In some conditions, platelets were pretreated with BI-1002494 (10µM) before addition of hemin. Following 3-hour incubation at 37 °C, cells were washed, fixed and stained with Sytox green for 5 min and

DNA stain imaged using EVOS microscopy. DNA area coverage of 5 separate fields/condition was measured using imageJ.

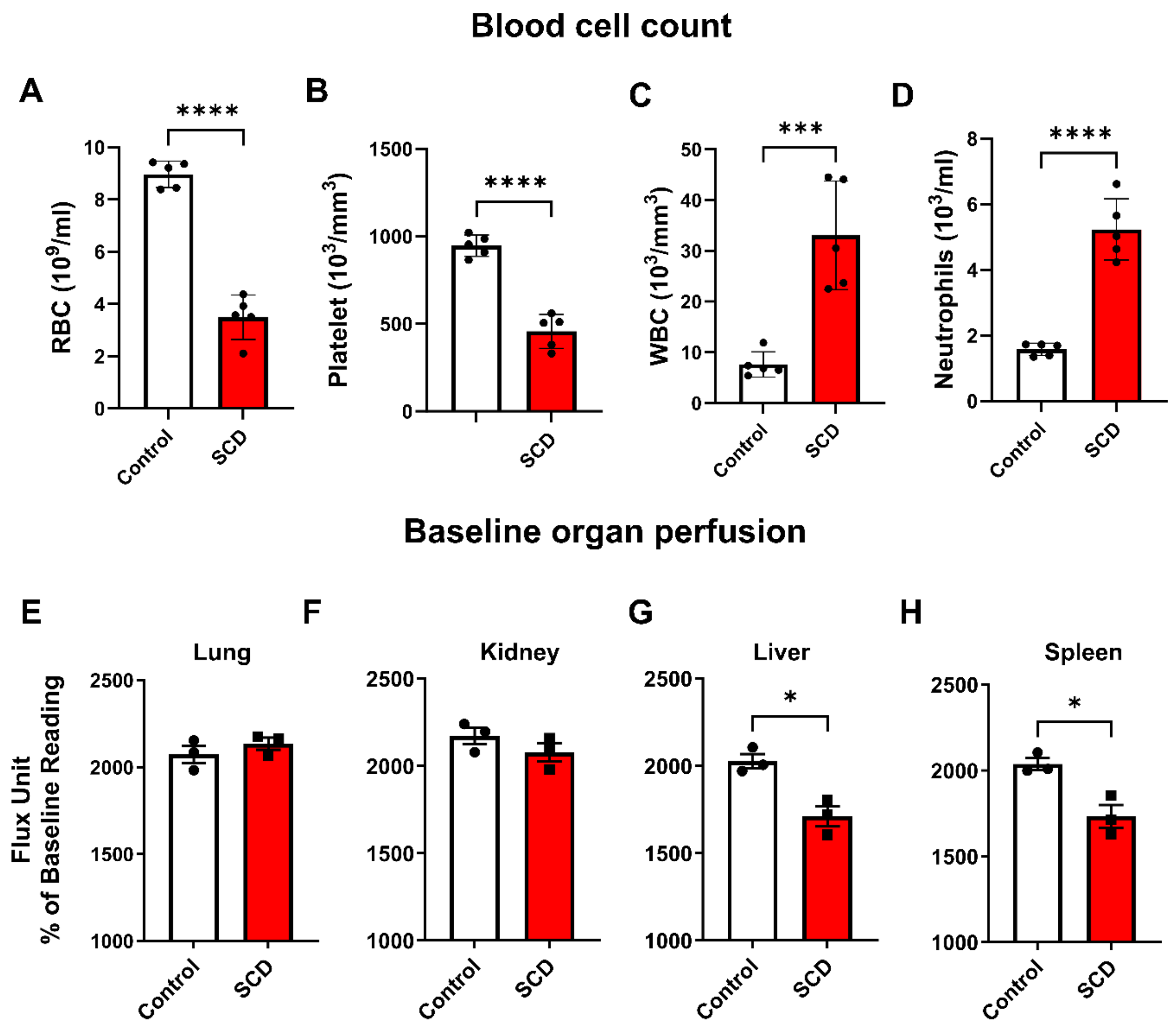


**Supplementary Figure 1: Blood cell count and organ perfusion at baseline in control non-sickle and SCD mice.** (A-D) Blood was collected from mice prior to hemin (at baseline). (A) platelet, (B) RBC, (C) WBC and (D) neutrophils in the blood were measured using an automated hematology analyzer. (E-H) Lung, kidney, liver and spleen perfusion was measured by laser speckly contrast imaging. Quantitative analysis of flux unit readings as a percentage of baseline values obtained by LSCI and the corresponding

AUC curve for the (E) lung, (F) kidney, (G) liver and (H) spleen. The statistical significance between 2 groups was analyzed using an unpaired t-test *p < 0.05, ****p<0.0001.

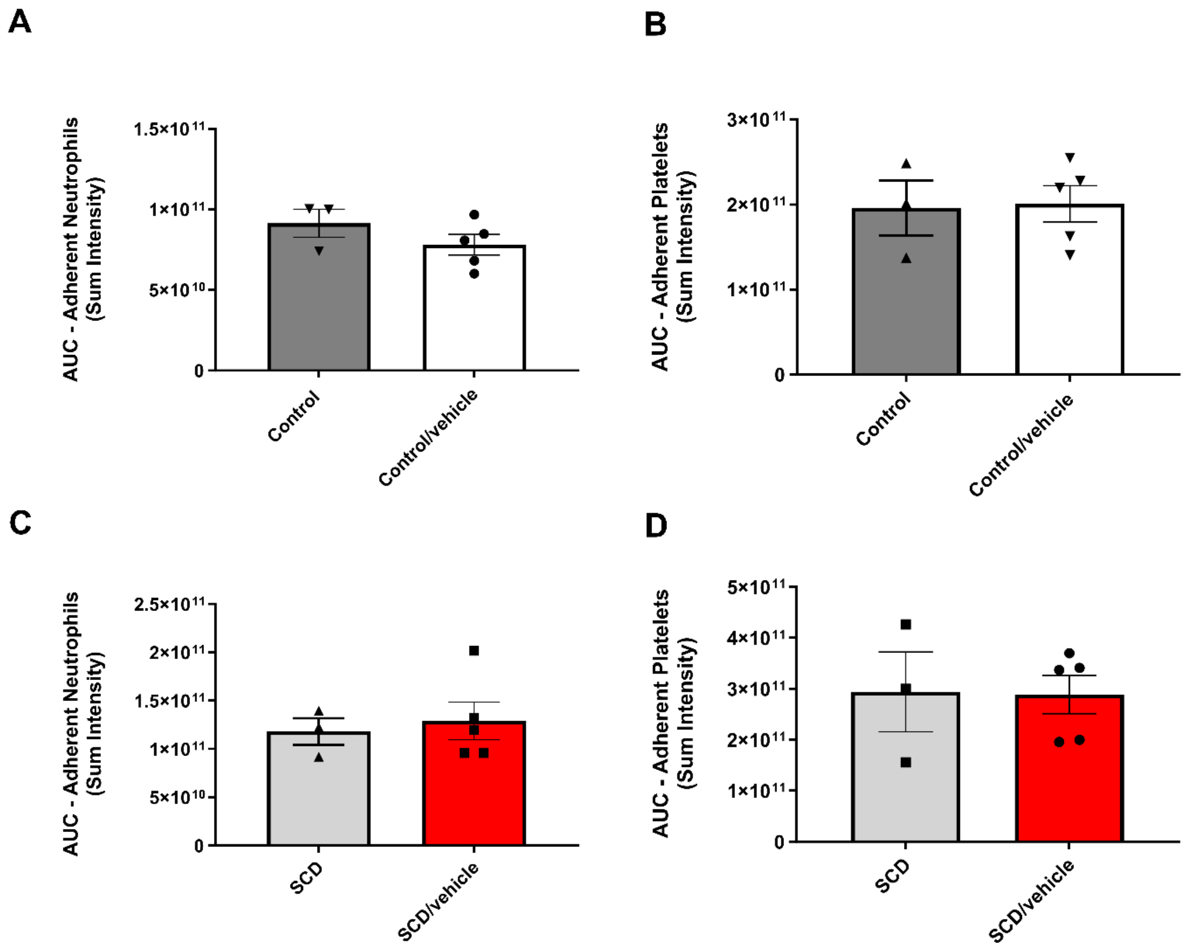


**Supplementary Figure 2: Pretreatment with vehicle does not alter platelet or neutrophil recruitment to the lung of control or SCD mice.** Hemin (20µmol/kg, I.V.) was injected to control and SCD mice. Vehicle (PBS-DMSO 10%) was injected intraperitoneally to mice 30 min prior to hemin injection. (A, C) Area under the curve (AUC) analysis over a time course of 60 min for neutrophils (Gr-1). (B, D) Area under the curve (AUC) analysis over a time course of 60 min for platelets (GPIbβ). The statistical significance between 2 groups was analyzed using an unpaired t-test.

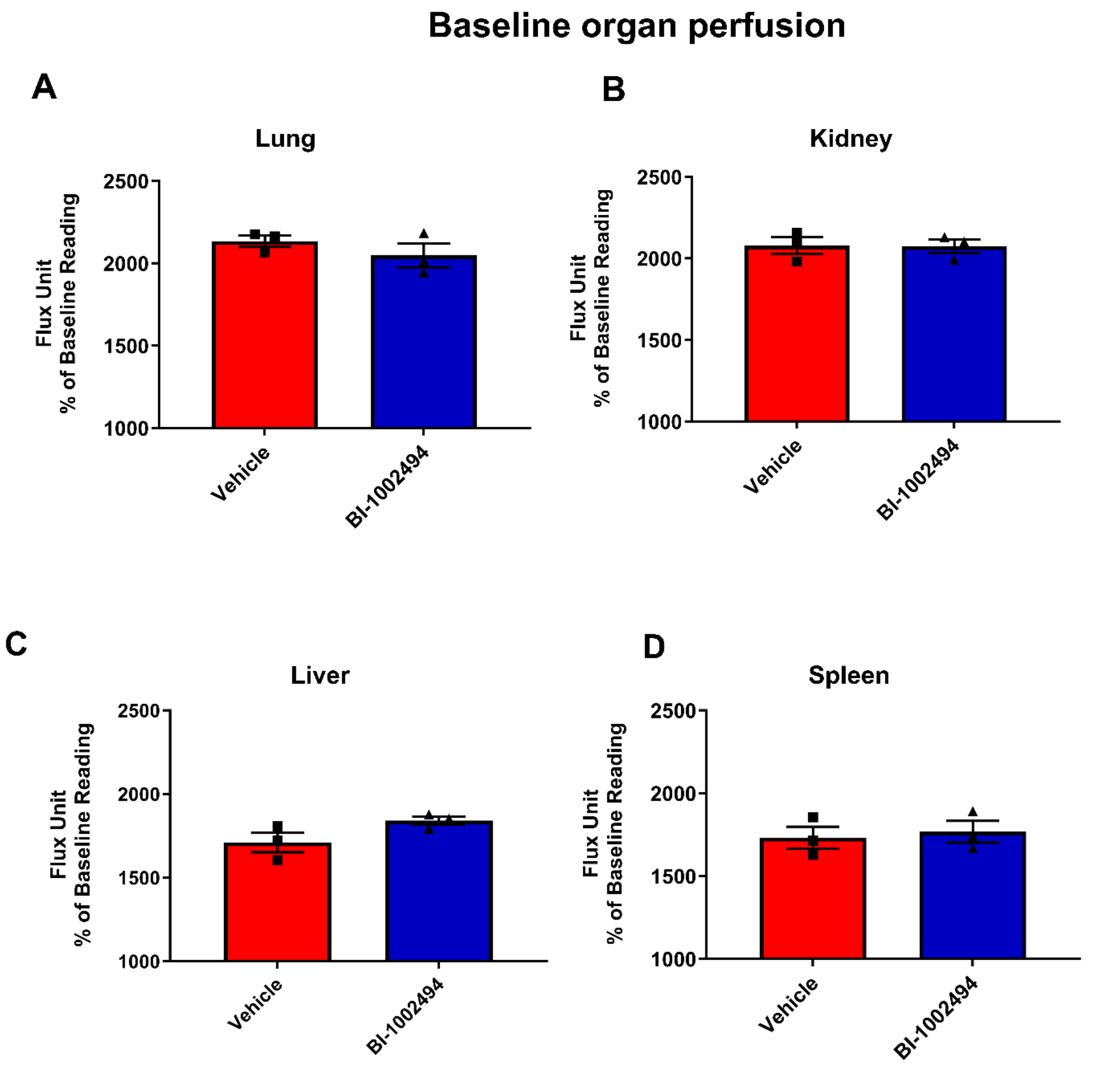


**Supplementary Figure 3: Syk inhibitor BI-1002494 pretreatment does not alter basal organ perfusion at baseline.** BI-1002494 (20mg/kg) or vehicle were injected in SCD mice via intraperitoneal route 30min before the injection of hemin (20µmol/kg, I.V.). Quantitative analysis of flux unit readings as a percentage of baseline values obtained by LSCI and the corresponding AUC curve for the (A) lung, (B) kidney, (C) liver and (D). The statistical significance between 2 groups was analyzed using an unpaired t-test.

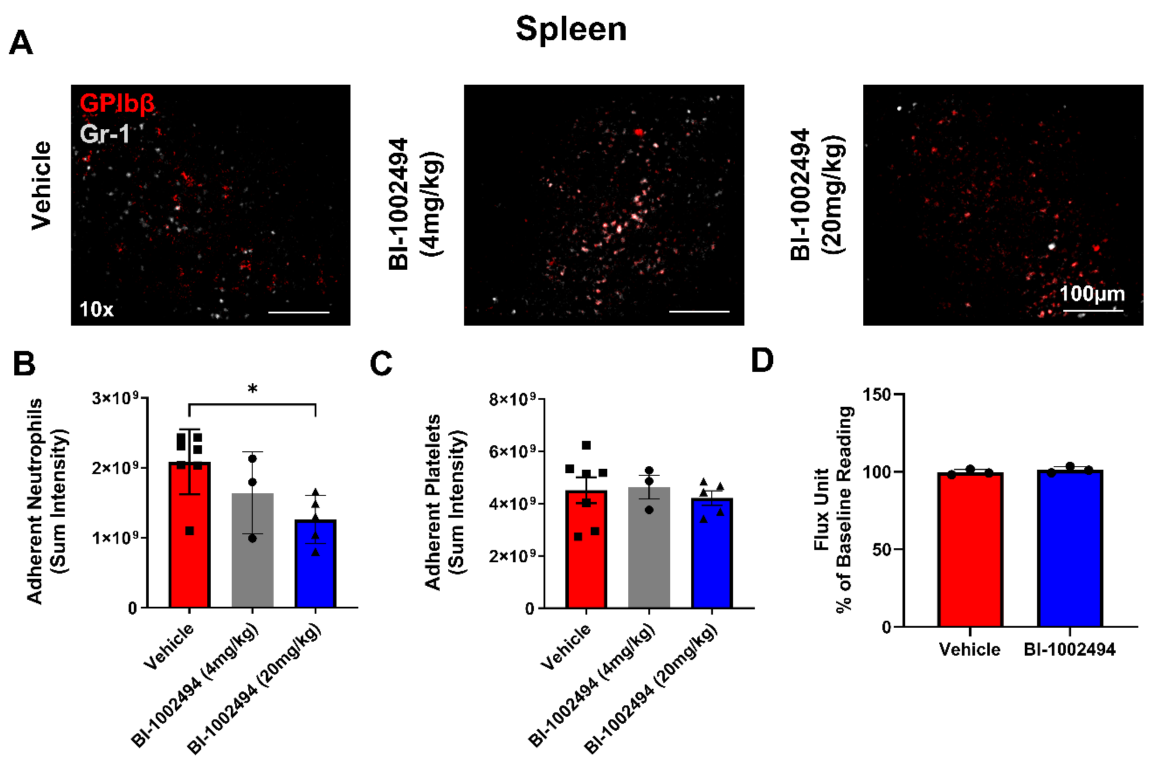


**Supplementary Figure 4: BI-1002494 reduces neutrophil recruitment to the spleen without alteration in perfusion.** BI-1002494 (4 or 20mg/kg) or vehicle were injected in SCD mice via intraperitoneal route 30min before the injection of hemin (20µmol/kg, I.V.) for 1h. (A) Representative intravital images of the breathing lungs showing adherent neutrophils (Gr-1; grey) and platelets (GPIbβ; red) in the splenic vasculature at 60 min post-hemin at 10x magnification. (B) Quantitative analysis of the intravital data for total adherent neutrophils and (C) platelets before injection of hemin and following 1h following hemin injection shown as the area under the curve (AUC). (D) Quantitative analysis of flux unit readings as a percentage of baseline values obtained by LSCI and the corresponding AUC curve. The statistical significance between multiple groups analyzed using Kruskal-Wallis test with multiple comparisons and plotted compared to control $*p < 0.05$.

**Video 1: Platelet and neutrophil recruitment to the lung following hemin injection in control non-sickle mice.**

**Video 2: Platelet and neutrophil recruitment to the lung following hemin injection in control SCD mice.**

**Video 3: Platelet and neutrophil recruitment to the lung following hemin injection in control SCD mice pretreated with BI-1002494 (20 mg/kg).**
